# Cure of experimental *Trypanosoma vivax* infection with a single dose of an unmodified antibody-based drug targeting the invariant flagellum cell surface protein IFX

**DOI:** 10.1101/2025.07.08.663690

**Authors:** Delphine Autheman, Cristina Viola, Georgina Rhodes, Simon Clare, Cordelia Brandt, Katherine Harcourt, Gavin J. Wright

## Abstract

Animal African trypanosomiasis (AAT) is an infectious wasting disease of economically important livestock caused by *Trypanosoma* spp. parasites. The disease is primarily caused by two species: *T. congolense* and *T. vivax* which are endemic in many African countries. AAT is managed by therapeutic and prophylactic drugs; however, resistance is now widely reported and the development of new drugs has been impeded due to a chronic lack of investment. Recently, we identified an invariant flagellar-associated cell surface protein (IFX) that could elicit protective immune responses when used as a vaccine against *T. vivax*. We showed that a complement-recruiting anti-IFX monoclonal antibody can prevent infection when used prophylactically. Here, we show that this same unmodified antibody can be used to cure *T. vivax* infections in a murine experimental model. Importantly, we show that infections can be treated with a single dose and demonstrate full cure by the lack of detectable parasites in peripheral tissues even after immunosuppression. Using structural modelling and site-directed mutagenesis, we localise the protective antibody epitope thereby identifying targetable regions on IFX to improve vaccine design. Together, these findings validate IFX as both a prophylactic and curative drug target that could be useful in the management of AAT.

**Importance:** *Trypanosoma vivax* is a parasite that causes animal African trypanosomiasis (AAT), a chronic wasting disease that infects economically-important livestock animals which is a particular problem in African countries south of the Sahara. The impact of this disease is significant: it is responsible for over 3 million cattle deaths and an estimated $4.5 billion of annual lost productivity. There is a desperate need to develop new control measures because resistance is now widely reported to the drugs commonly used to treat this infection. We show here that a single dose of an unmodified monoclonal antibody that recognises IFX - a parasite cell surface protein localised to the flagellum - is sufficient to cure an established *T. vivax* infection with no parasite reservoirs detectable in peripheral tissues. Our finding validates IFX as a new drug target and provides a rationale route to the development of new drugs to target AAT.

## Introduction

The livelihoods of millions of people living in Africa are at risk due to infectious diseases that affect the health of their livestock which provide them with essential food, milk, clothing, manure and draught power. One major livestock disease is animal African trypanosomiasis (AAT) which is caused by blood-dwelling parasites of the genus *Trypanosoma* that affect many important farm animals including sheep, pigs, and especially cattle (1). The impact of this disease is significant: over 3 million cattle die from this disease annually with a direct economic cost to the African economy amounting to many hundreds of millions of dollars (2). This disease therefore represents a major constraint on the economic development of the mainly agrarian societies in African countries south of the Sahara (3, 4).

Although several species of trypanosome parasite can cause this disease in animals, the two most prevalent are *T. congolense* and *T. vivax*. The disease is currently managed by controlling the tsetse fly vector population and the use of trypanocidal drugs (5). The most commonly used drugs for therapy and prophylaxis are diminazene aceturate and isometamidium chloride: both were developed over seventy years ago in the 1950s (6, 7). These drugs can cause toxic side effects and when inappropriately administered, for example by underdosing, has led to the emergence of drug resistance which has now been reported in 21 out of the 37 countries in which the disease is endemic (8). The development of new trypanocides for livestock has lacked sufficient investment because there is a poor economic case: new drugs remain expensive to develop, and a price point that would provide a satisfactory return for developers could not be met by the primary customers who are largely subsistence farmers. Despite this, there has been some encouraging recent progress achieved by the repurposing of drugs developed to treat human African trypanosomiasis (9–11).

Another control tool would be the development of an effective vaccine, but none yet exist for AAT (12). Vaccination has generally been considered a low value option for this disease because the parasite has evolved sophisticated immunoregulatory mechanisms such as antigenic variation (13), the ability to rapidly clear surface bound host antibodies (14), and disruption of B-cell development and maintenance (15, 16) all of which enable trypanosomes to thrive in the blood of their infected host. We have previously shown that it is possible to induce protective immunity in an experimental model of *T. vivax* infection by eliciting an antibody response to an invariant flagellum-associated antigen called IFX prior to challenge by the parasite (17). In our experiments to determine the immunological mechanisms of protection, we identified a small panel of monoclonal antibodies, some of which could confer protection to *T. vivax* infection by passive transfer. While the original hybridoma-derived IgG1-isotype antibodies conferred only modest protection, reformatting the isotype of one antibody called 8E12 as an immune-effector-recruiting IgG2a isotype could elicit very potent protection when delivered prophylactically. We additionally demonstrated that by mutating the C1q and FcR binding sites in the Fc region of the antibody, we showed that complement recruitment was the major protective mechanism followed by FcR binding, and lastly direct opsonization of the IFX antigen (17).

The passive transfer of immune serum from convalescent patients to treat and prevent infectious diseases has a long history of success (18). This approach, however, poses challenges in the clinic because it requires a sufficiently large number of donors, often varies in its efficacy, and carries the risk of unintentionally transferring infections. Success in treating a range of diseases from cancer to autoimmune disease using monoclonal antibody therapeutics has driven biotechnological progress that have made the isolation and large-scale production of antibody-based drugs rapid and cost effective (19). Increasingly, this class of drugs has been developed for the treatment and prevention of infectious diseases (20). For example, several antibody-based drugs targeting the SARS-CoV-2 spike protein were developed within a few months during the COVID-19 pandemic including bamlanivimab, sotrovimab, and regdanvimab (21, 22). A major advantage of these drugs are their long pharmacokinetics, lack of toxicity, and generalised production methods (23). As well as treating viral infections, this strategy has also been shown to work as both therapy and prophylaxis for bacterial infections (22, 24). Importantly, antibody-based therapies have also been used to treat and prevent parasitic infections. This includes antibodies that target the circumsporozoite protein of sporozoite-stage *Plasmodium falciparum* parasites which showed dose-dependent prevention of infection in Malian patients over a period of six months (25, 26). Similarly, experimental *Trypanosoma brucei* infections could be cured with a single dose of an antibody-drug conjugate targeting the haptoglobin-haemoglobin receptor (27).

Here, we investigate the possibility of using a monoclonal antibody targeting the *T. vivax* vaccine antigen IFX in a drug-like manner. We use structural modelling to identify the epitope of the 8E12 antibody and show using a murine infection model that a single dose of the unmodified antibody can cure an established *T. vivax* infection.

## Results

### Structural modelling identified the location of the 8E12 epitope on IFX

To explore the possibility of using the IFX protein as a potential drug target, we first characterised the anti-IFX 8E12 monoclonal antibody that could elicit passive protection to *T. vivax* infections in more detail. We sequenced the rearranged light and heavy chains of the 8E12 antibody which had been amplified from a hybridoma to create a single plasmid (28). Sequencing of this plasmid revealed kappa light chain usage and identified the amino acids forming the antibody paratope (Supplementary Figure 1). In our original study, we determined the 8E12 binding affinity as only moderate (*K*_D_=14nM), and mapped the 8E12 epitope to the N-terminal 168 amino acids of IFX by expressing fragments of the IFX ectodomain (17). To characterize the 8E12 epitope further, we first asked if the epitope was conformational by denaturing the IFX protein by heat treatment and reduction. We observed that while some of the binding activity was lost upon both heat treatment and protein reduction, not all binding activity was lost indicating that the 8E12 epitope was at least partially dependent on protein folding (Fig. 1a). There is no experimental structural information on IFX, but recent advances in protein structure (29) and binding interface prediction (30) presented the opportunity to model both the IFX structure and predict the 8E12 epitope. Structural modelling suggests that the extracellular region of IFX adopts a dumbbell shape composed of two groups of alpha-helices joined by a long central helix (Fig. 1b). By using the sequence of IFX and the light and heavy chains of the 8E12 antibody, we used Alphafold3 to predict the location of possible epitopes. Of the five most highly-ranked models, all converged on a region of IFX at the tip of one of the helical clusters (Fig. 1b). We were reassured by the convergence of these predictions with the experimentally known N-terminal location of the epitope, and the lack of overt steric clashes with the location of N-linked glycans.

**Figure 1.**
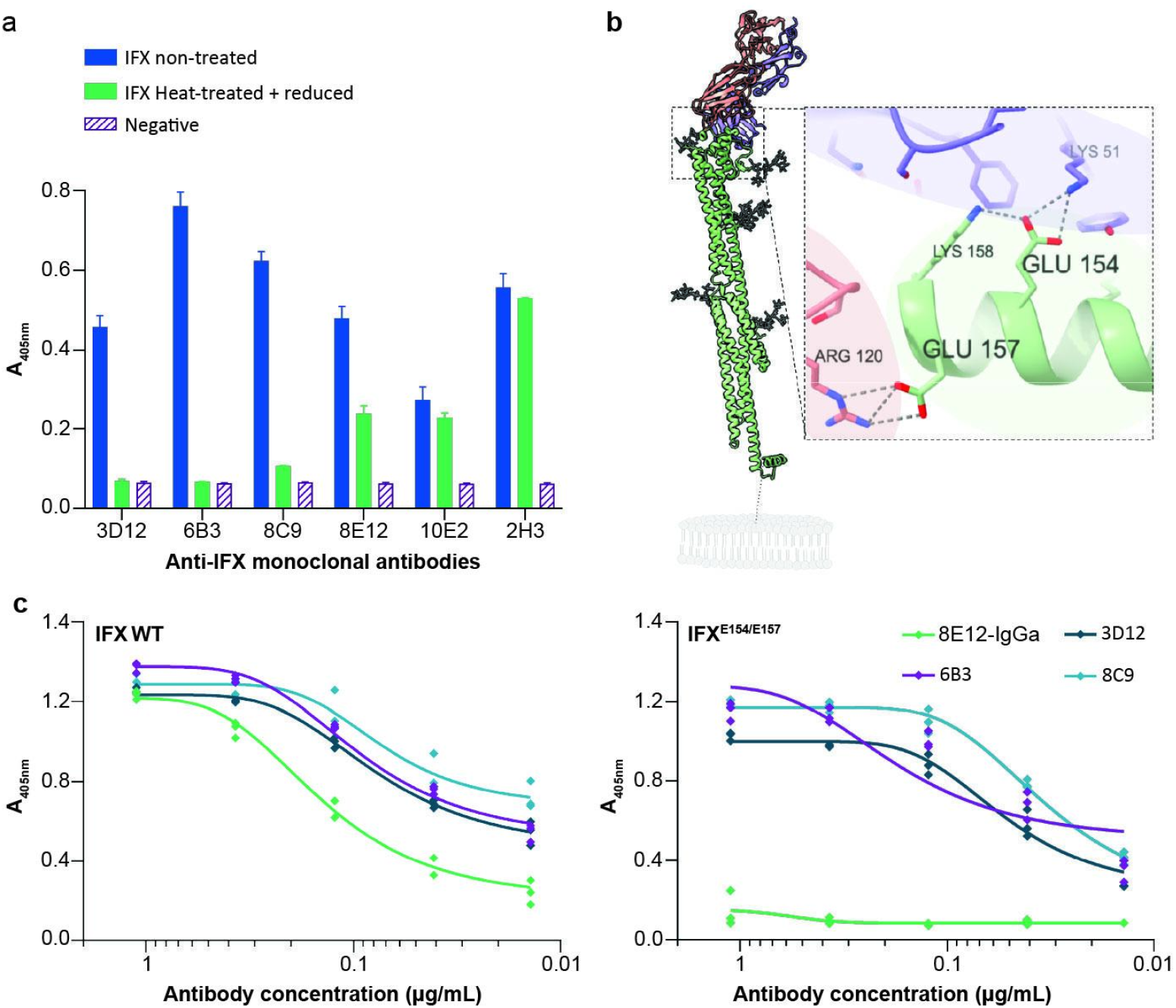
Identifying the location of the anti-IFX 8E12 monoclonal antibody epitope by structural modelling and site-directed mutagenesis. **(a)** Biochemical characterisation of anti-IFX monoclonal antibody epitopes. The monobiotinylated IFX ectodomain was immobilised in wells of a streptavidin-coated microtitre plate without treatment or after heat treatment with reduction. Immunoreactivity of antibodies 3D12, 6B3 and 8C9 was completely abrogated in the denatured protein showing they recognised a conformational epitope; immunoreactivity to 2H3 and 10E2 was retained upon denaturation demonstrating these antibodies recognized a non-conformational epitope. Immunoreactivity to 8E12 was only partly ablated by heat treatment and protein reduction demonstrating the epitope was only partly conformational. Bars represent means ± SD *n* = 3. **(b)** Structural model of the IFX-8E12 Fab complex. The predicted structure of the extracellular region of IFX (green), including the location of modelled potential N-linked glycans (gray sticks) in complex with the 8E12 antibody Fab fragment showing light chain (red) and heavy chain (blue). Inset: atomic details of the 8E12 epitope highlighting the position of glutamic acid residues at positions 154 and 157. The approximate location of the parasite plasma membrane phospholipids are shown in gray. **(c)** Identification of the epitope recognised by the 8E12 antibody. The entire extracellular region of IFX was expressed as either its wild-type form (left panel) or the E154K / E157K mutant (right panel) as a soluble biotinylated protein and immobilised in wells of a streptavidin-coated plate. Titrations of the indicated anti-IFX antibodies were tested for direct binding to the IFX proteins by ELISA. While both wild-type and mutant IFX bound antibodies 3D12, 6B3 and 8C9, the 8E12 antibody bound only the wild-type form.

To experimentally validate these predictions, we identified two solvent-exposed glutamic acid residues (E154 and E157 of TvY486_0807240) which contributed hydrogen bonds in the 8E12 co-complex structure prediction and mutated these to lysine. We showed that E154 and E157 contributed to the 8E12 epitope by showing that the mutant IFX protein was unable to bind the 8E12 antibody (Fig. 1c). Importantly, the mutant IFX retained the ability to bind other anti-IFX monoclonal antibodies demonstrating the mutations did not cause significant mis-folding of the IFX ectodomain (Fig. 1c). Together, these data identify the 8E12 epitope to a membrane-distal tip of a helical cluster on the IFX ectodomain.

### Repeated doses of 8E12 mAb can cure an experimental *T. vivax* infection

In our initial experiments, 8E12 was used prophylactically to prevent infection by administering the 8E12 antibody the day before, on the day, and the day after infection with *T. vivax* (17). To be used in a drug-like manner, the antibody must be able to clear an established infection. First, a bulk preparation of the 8E12 monoclonal antibody was produced by transfecting HEK293 cells and purified using immobilised protein G. The antibody was shown to be highly active by quantifying binding to recombinant IFX by ELISA and over 95% pure by resolving the preparation on a protein gel (Supplementary Figure 2). Next, we infected mice with a transgenic form of *T. vivax* constitutively expressing the firefly luciferase, and waited until four days post infection where a patent infection was apparent, and could be quantified using bioluminescent imaging. At day five postinfection, the average bioluminescence was 4.1×10^7^ photon s^-1^ which corresponds to a parasitaemia of ∼1×10^6^ parasites per mL of blood, a value that is towards the highest amount detected in infected cattle (31). From day five, we administered a total of three daily doses of 50, 100, 200, or 400 micrograms of purified 8E12 IgG2a-formatted antibody and continued to measure parasitaemia each day (Fig. 2a). We observed that the lowest (50μg) doses, while slowing the multiplication of the parasite, was not able to clear the infection and all five animals were removed from the study by day nine (Fig. 2b, c). While 100μg doses could reduce parasitemia in most animals to very low levels by day eight, there was a recrudescence of the infection in each animal such that all had to be removed from the study by day 15 (Fig. 2b, c). At 200μg, parasites were reduced below the limits of detection by day seven (Fig. 2b), but then showed recrudescence of this infection that was highly variable between individuals. Interestingly, two of the five treated animals showed the ability to control the infection after day 20 which significantly exceeds the measured *in vivo* half life for an IgG2a-isotype antibody which has been measured at 6 to 8 days (32) suggesting that the animals had acquired some immunity. All animals receiving 400μg doses showed no evidence of recrudescence with no detectable parasites from day seven until beyond day 60.

**Figure 2.**
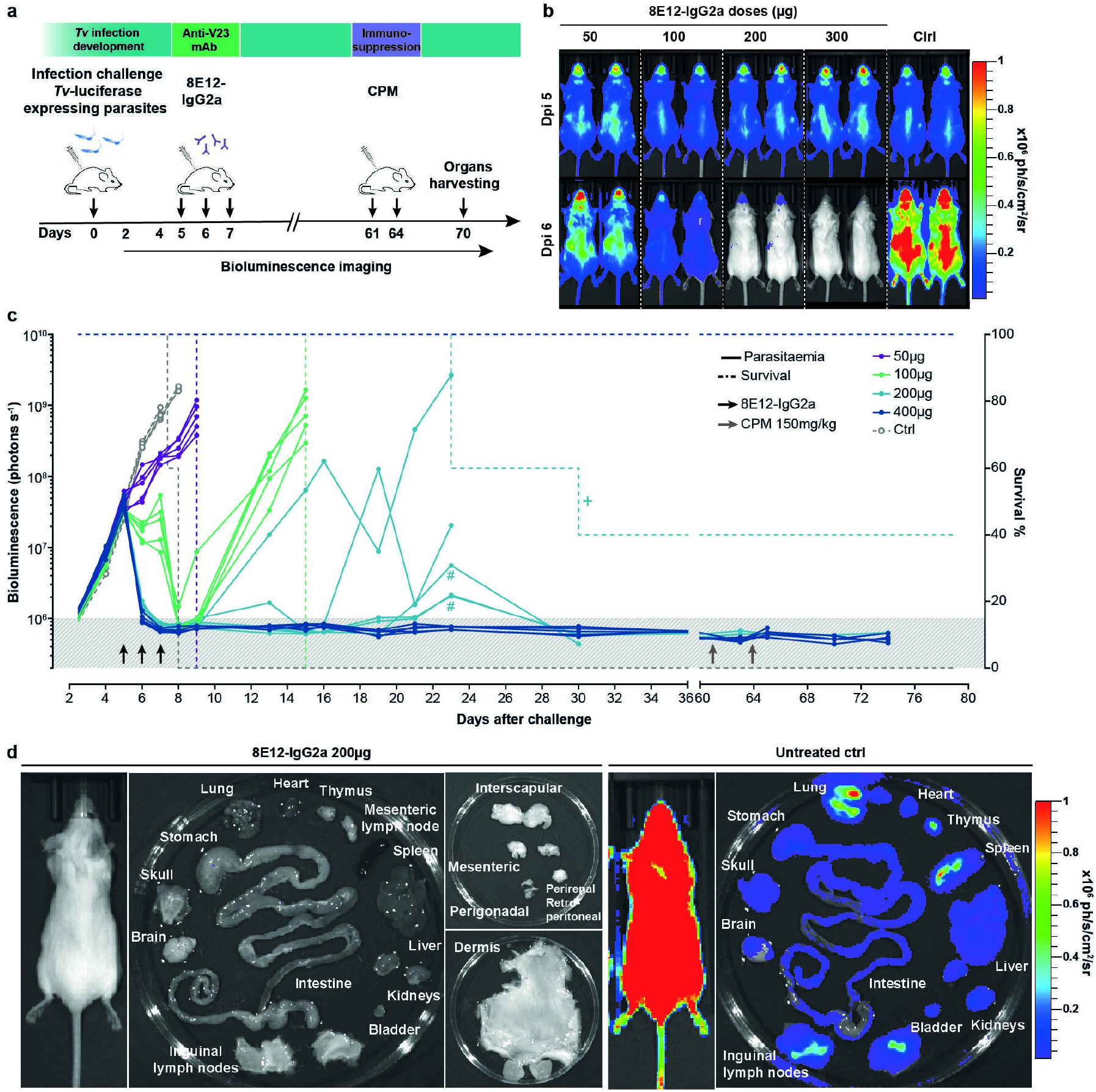
Cure of an experimental *T. vivax* infection with the anti-IFX 8E12-IgG2a-formatted monoclonal antibody. (**a**) Schematic showing the experimental plan. Groups of five animals were infected with luciferase-expressing transgenic *T. vivax* parasites on day 0. Varying doses of the 8E12 monoclonal antibody were administered daily on days 5, 6, and 7 and parasitaemia was quantified by bioluminescent imaging. Animals were immunosuppressed by two doses of cyclophosphamide (CPM). (**b**) Images of mice infected with bioluminescent *T. vivax* immediately before and 24 hours after the last administration of the indicated doses of the 8E12 antibody. (**c**) Longitudinal analysis of individual infected animals with the indicated doses of 8E12 relative to an isotype-matched control antibody (Ctrl). Bioluminescence (solid lines) and survival (dashed lines) are plotted for each experimental group. Background level of bioluminescence is indicated by grey shading. Crosses indicate where animals were removed from the study for welfare reasons that are thought to be unrelated to the infection. Black and grey arrows represent doses of 8E12 antibody and injection of the cyclophosphamide immunosuppressant respectively. Hash symbols represent potential bioluminescence signal leakage from a heavily infected mouse to an adjacent mouse with a lower bioluminescence signal during image acquisition. (**d**) Lack of residual parasites in peripheral tissues of treated and immunosuppressed animals. No bioluminescent parasites were detected in the named tissues of a *T. vivax*-infected mouse that was administered with three 200μg doses of the 8E12 antibody and subsequently treated with an immunosuppressant (left panels), relative to a control infected animal (right panels).

It is known that trypanosomes can reside in peripheral tissues such as the brain and adipose tissue which may have limited accessibility to antibody-based drugs. To ensure that there were no residual parasites that were being suppressed by the acquisition of immunity during the course of the experiment, we immunosuppressed apparently cured animals with two doses of cyclophosphamide at days 61 and 64, and observed the animals for a further fifteen days. In no case did we observe any recrudescent infections. To further ensure that no parasite reservoirs remained, we dissected out tissues of cured and immunosuppressed mice and subjected them to sensitive *in vivo* imaging. Again, we observed no parasite reservoirs in cured animals (Fig. 2d).

### Cure of an experimental *T. vivax* infection with a single dose of 8E12 antibody

Cattle are the main target species for drugs that treat animal African trypanosomiasis and because they usually are left to roam freely around the bush and handling facilities are often rudimentary, being able to treat the disease with a single dose would be a major advantage. To determine whether animals could be cured by a single dose, we again infected groups of mice with *T. vivax* before administering a single dose of either 300, 400 or 600μg of the 8E12-IgG2a-formatted antibody (Fig. 3a). To determine how rapidly parasites were killed, we imaged all animals at 4 and 24 hours after the time of treatment. Consistent with our earlier finding that complement plays the major role in the protective immunological mechanism (17), we observed a dramatic reduction in parasitaemia at four hours which progressed to below the limit of detection by 24 hours (Fig. 3a, b). There was a recrudescence of the infection in all animals receiving the 300μg dose such that all animals were removed from the study by day 14. Two out of the five animals receiving the 400μg dose were apparently cured of the infection, and a further two animals showed a drastic reduction in parasitaemia by day 14 suggesting acquisition of adaptive immunity (Fig. 3a). All animals receiving a single 600μg dose were apparently sterilely cured. To demonstrate that there were no remaining parasites below the limits of detection, we immunosuppressed animals on day 48, and showed no recrudescence of the infection. Again, we dissected tissues of remaining immunosuppressed animals on day 57 and showed no evidence of residual parasites in peripheral tissues (Fig. 3c).

**Figure 3.**
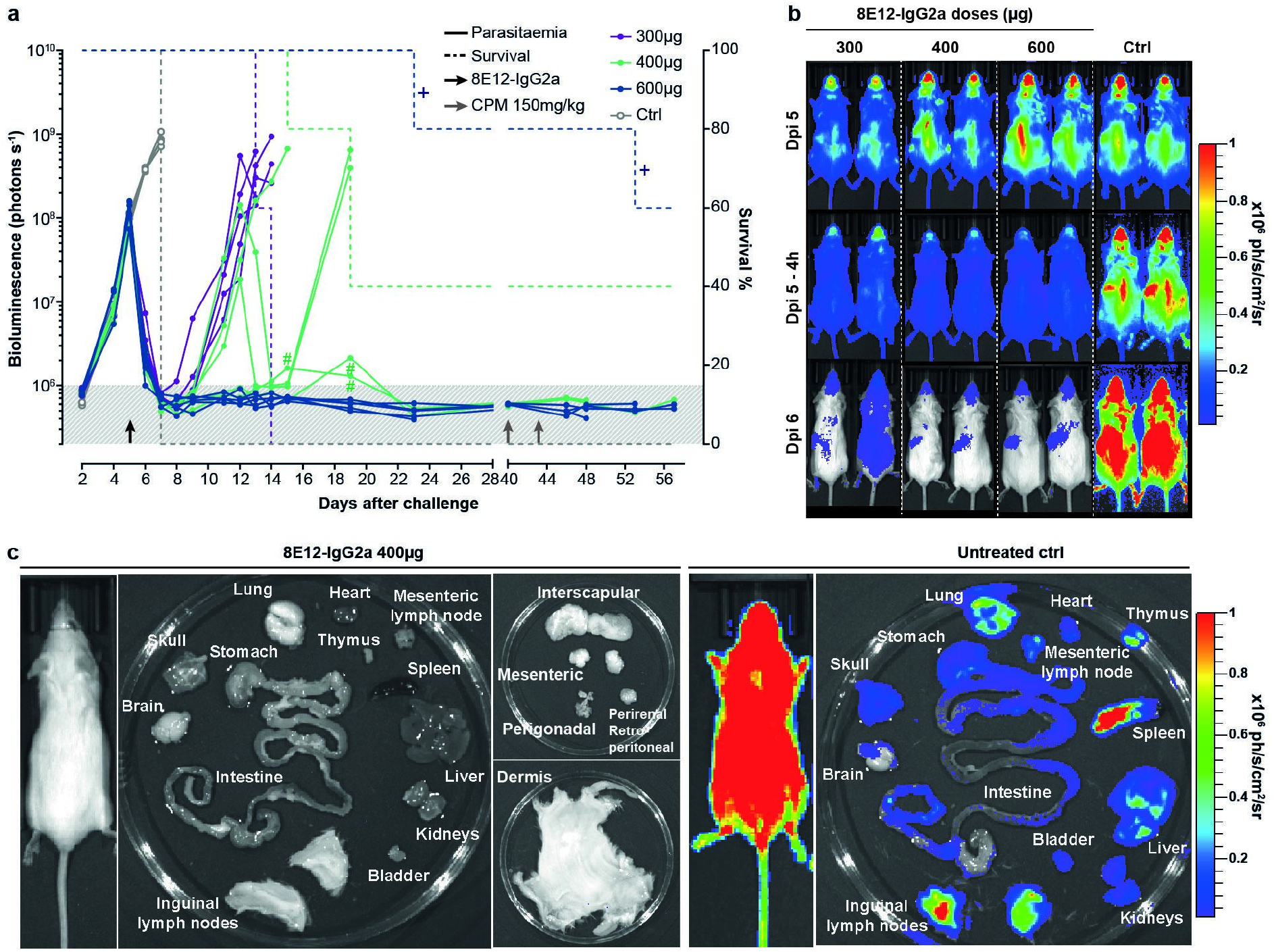
Rapid cure of an experimental *T. vivax* infection with a single dose of anti-IFX 8E12-IgG2a-formatted monoclonal antibody. (**a**) Longitudinal analysis of individual infected animals with the indicated single doses of 8E12 relative to an isotype-matched control antibody (Ctrl). Bioluminescence (solid lines) and survival (dashed lines) are plotted for each experimental group. Background level of bioluminescence is indicated by grey shading. Crosses indicate where animals were removed from the study for welfare reasons thought to be unrelated to the infection. Black and grey arrows represent the single dose of 8E12 antibody and injection of cyclophosphamide (CPM) immunosuppressant respectively. Hash symbols represent potential bioluminescence signal leakage from a heavily infected mouse to an adjacent mouse with a lower bioluminescence signal during image acquisition. (**b**) Dorsal views of mice infected with bioluminescent *T. vivax* at 0 (Dpi 5), 4 (Dpi 5 - 4h) and 24 (Dpi 6) hours after the administration of the indicated doses of the 8E12 antibody showing rapid cure of infection. (**c**) Lack of residual parasites in peripheral tissues of treated and immunosuppressed animals. No bioluminescent parasites were detected in the named tissues of a *T. vivax*-infected mouse that was administered with a single 400μg dose of the 8E12 antibody and subsequently treated with an immunosuppressant (left panels), relative to a control infected animal (right panels).

## Discussion

Here we have demonstrated that a single dose of a purified but otherwise unmodified monoclonal antibody can cure an established parasitic infection. This leads to the wider question of whether this general strategy of using antibody-based drugs could be a realistic treatment option for parasitic diseases of livestock. It is well known that the passive transfer of polyclonal immune serum from convalescent patients or immunised animals can be an effective treatment for infectious diseases and neutralising toxins in humans (33–35). This approach is unlikely to be a practical option for an endemic disease of livestock because with an estimated 350 million heads of cattle in Africa, this number is impractically large (36). The general view is that therapeutic monoclonal antibodies are too expensive for use in livestock, but recent technological developments in isolating and producing large amounts of monoclonal antibodies driven by the multi-billion dollar therapeutic antibody industry has made these drugs much more affordable. The adoption of generalised production methods and economies of scale mean that gram quantities of purified antibody can be produced for as little as $10 (37), and it is likely that these costs will continue to decrease. These factors have spurred the use of therapeutic antibodies to treat infectious diseases either by targeting susceptible pathogen proteins or even host proteins (38–40).

Clearly, there would be some challenges for the use of such a strategy for the treatment of AAT. Firstly, this disease is endemic in developing countries where a reliable cold chain to ensure the protein-based product retains activity cannot be assumed. Advances in the design of heat-stable affinity scaffolds, however, provide confidence that this problem could be overcome (41, 42). Also, antibody accessible cell surface proteins are often polymorphic so that specific antibody-based reagents will almost certainly be species-specific meaning that a single drug would be unlikely to be effective against different species of trypanosome. IFX, for example, is only found in *T. vivax* with no easily identifiable orthologues in other sequenced species (17). Because *T. congolense* and *T. vivax* are co-endemic in Africa, this would require the development of two different drugs, although a drug targeting just *T. vivax* would be useful in regions such as South America which has a large cattle industry and where *T. vivax* is the primary cause of AAT.

Administration of antibody doses that were not sufficient for cure typically resulted in a rapid reduction in parasitaemia but that resulted in later recrudescence of the infection. In some animals, these later waves of parasitaemia could be controlled, even more than two weeks after the last antibody dose. This suggests that the adaptive immune system had a role in controlling the infection at this stage because the half life of circulating antibodies is typically around a week in mice. This may suggest that *T. vivax* infections do not significantly disable the adaptive immune system in the way that has been reported for *T. brucei* infections in mice (16, 43).

Targeting infectious diseases with antibody-based drugs does have some advantages. The antibody described here can be used in both a curative and prophylactic manner. This would be important for cattle farmers in Africa who generally lack adequate animal holding facilities and so a single dose drug that would cure current infections and provide lasting protection would be valuable and more practical to administer. Antibodies have excellent pharmacokinetics with circulating half-lives of several weeks due to their relatively large size and active recycling mechanisms mediated by the neonatal Fc receptor (FcRn) (44).

Engineering of the antibody constant regions that interact with FcRn have enabled circulating half lives to be significantly extended (45, 46). For use in cattle, it would be important to bovinise the antibody so that it effectively recruits immune effectors and does not elicit host anti-drug immune responses. The recent sequencing, annotation, and functional characterisation of the bovine antibody locus now provides the necessary information to achieve this (47).

Motivated by the use of antibody-drug conjugates (ADCs) that have been used to treat cancer, research from MacGregor and colleagues showed that an experimental *T. brucei* infection could be cured with a single dose of an antibody targeting the haptoglobin-haemoglobin receptor conjugated to the toxin pyrrolobenzodiazepine (27). While initially conceived as a curative drug for human use, a similar ADCs strategy could increase the potency of similar drugs for use in livestock. Here though, there may be a conflict between trying to increase the *in vivo* half-life of the drug to provide prophylactic cover, and the need to ensure there is no long-lasting drug toxicity remaining in livestock products such as the milk and meat which would be undesirable for consumers. In summary, the work described here lends further support for the use of antibody-based drugs to treat livestock infectious diseases and identifies IFX as a new curative and preventative drug target for animal African trypanosomiasis.

## Materials and Methods

### Mouse strains and ethical approvals

All mouse experiments were performed under UK Home Office governmental regulations (project licence numbers PD3DA8D1F and P98FFE489) and European directive 2010/63/EU. Research was ethically approved by the Sanger Institute Animal Welfare and Ethical Review Board. Mice were maintained under a 12-h light/dark cycle at a temperature of 19–24 °C and humidity between 40 and 65%. The mice used in this study were 6–14-week-old female *Mus musculus* strain BALB/c, which were obtained from a breeding colony at the Research Support Facility, Wellcome Sanger Institute.

### Statistical analysis

The experiments were not randomized, and investigators were not blinded to allocation during experiments and outcome assessment, although animal dosing and parasite quantification were performed by independent researchers.

### Cell lines and antibodies

Recombinant proteins and antibodies used in this study were expressed in HEK293-6E cells provided by Y. Durocher. Cell lines were not authenticated but were tested for mycoplasma. The 8E12 antibody was cloned and expressed recombinantly as a mouse IgG2a isotype as described (17, 28). The mouse IgG2a isotype control antibody was C1.18.4, BioXcell Cat. No. BE0085. The other anti-IFX antibodies, 3D12, 6B3, 8C9, 10E2 and 2H3 were produced and purified by hybridoma supernatant as described (17). The antibody used for protein quantification in ELISAs was a mouse monoclonal anti-His (His-Tag monoclonal antibody, 70796, EMD-Millipore). The secondary antibody used was a goat anti-mouse alkaline phosphatase conjugated secondary (A3562, Sigma-Aldrich).

### Recombinant protein and antibody expression and purification

The recombinant soluble IFX ectodomain and controls were expressed in HEK293-6E cells as described (17). The IFX E154K/E157K mutant was produced using gene synthesis by Twist Biosciences. Proteins were expressed as enzymatically monobiotinylated proteins by co-transfection with a secreted version of the protein biotin ligase (BirA) as previously described (<u>Kerr et al. 2012</u>). Supernatants were collected five days after transfection, filtered and stored at 4°C until use. Proteins were purified using Ni^2+^ immobilized metal-ion affinity chromatography as described (48).

### ELISAs

To determine if the anti-IFX antibodies recognised a conformational epitope, the purified monobiotinylated ectodomain of IFX was denatured by heat treatment (90°C) in the presence of 10 millimoles of DTT for 10 minutes; once cooled 20 millimoles of iodoacetamide was added. The immunoreactivity of the 8E12 monoclonal antibody to the entire ectodomain of wild type IFX and the E154K/E157K mutant was quantified by ELISA. Purified ectodomains were normalised by dilution in PBS containing 0.2% Tween-20 (PBST) and 2% BSA and captured on a streptavidin-coated microtitre plate by incubating overnight at 4°C. Plates were washed two times with PBST, and once with PBS. Purified anti-IFX monoclonal antibodies were diluted in PBS/2% BSA, added to the ELISA plate and incubated overnight at 4°C. A 1 in 1000 dilution of an anti-6His antibody (EMD millipore) was used as a control to ensure ectodomains were captured. The plate was washed two times with PBST, and once with PBS, before incubating for one hour with a 1 in 5000 dilution of anti-mouse IgG secondary antibody conjugated to alkaline phosphatase (Sigma-Aldrich). After two further washes with PBST, and one wash with PBS, 70 μL of 1 mg/mL Sigma 104 phosphatase substrate was added, and substrate hydrolysis quantified at 405 nm using a plate reader (Spark, Tecan).

### Structural modelling

The three amino acid sequences encoding the light and heavy chains of the 8E12 Fab region and entire ectodomain of IFX (Uniprot: G0TZT9) were used as an input for AlphaFold3 for structural modeling. Models that exceeded the template modeling (pTM) and interface predicted template modeling (ipTM) scores of 0.5 and 0.8 respectively, and with the highest ranking predicted ‘Local Distance Difference Test’ pLDDT scores were selected for residue mutation.

### Quantification of *T. vivax* infections by bioluminescent *in vivo* imaging

Mice were infected with bloodstream forms of *T. vivax* parasites from the blood of an infected donor mouse at the peak of parasitaemia. Parasites were diluted in PBS and 20 mM D-glucose, quantified by microscopy, and used to infect mice by intravenous injection. The luciferase substrate D-luciferin (potassium salt, Source BioScience) was resuspended to 30 mg mL^−1^ in Dulbecco’s PBS (Hyclone), filter-sterilized (0.22 μm), and stored in aliquots until use at −20 °C. Luciferin was administered by intraperitoneal injection at 200 mg kg^−1^ ten minutes prior to imaging. Mice moved freely for three minutes before being anaesthetized (2.5% isoflurane) and placed in the imaging chamber where anaesthesia was maintained. The average background bioluminescence measurement was determined by luciferin administration in five uninfected female BALB/c mice and calculating the mean whole-body bioluminescence. This background value is indicated as a light grey shading on bioluminescence plots where appropriate. Long-term persistence of the parasites in different organs of infected mice was determined by imaging mice which were then euthanized with an overdose of anaesthetic. Mice were perfused with PBS until the perfusion fluid ran clear, the organs were dissected, and arranged on a Petri dish and bathed in PBS containing 20 mM glucose and 3.3 mg mL^−1^ luciferin. Emitted photons were acquired by a charge coupled device (CCD) camera (IVIS Spectrum Imaging System, Perkin Elmer). Regions of interest (ROIs) were drawn and total photons emitted from the image of each mouse were quantified using Living Image software version 4.7.4 (Xenogen), the results were expressed as the number of photons s^−1^. Bioluminescence values were exported and plotted in Prism GraphPad version 8.0.2, which was also used for testing statistical significance where needed. Persistence of potential parasite reservoirs in 8E12-IgG2a treated mice was assessed by immunosuppression. Mice received two doses of cyclophosphamide (150mg/kg, Cayman Chemical) three days apart using intraperitoneal injections and were monitored by imaging to assess parasite recrudescence.

## Acknowledgements

This research was funded by the Wellcome Trust (grant 206194) and the Gates Foundation (INV-005870). We thank Sanger Institute animal technicians for their support and Craig Mackenzie and Ben Luisi for preliminary work on anti-IFX monoclonal antibodies.

## Supplementary Figures

**Supplementary Figure 1.**
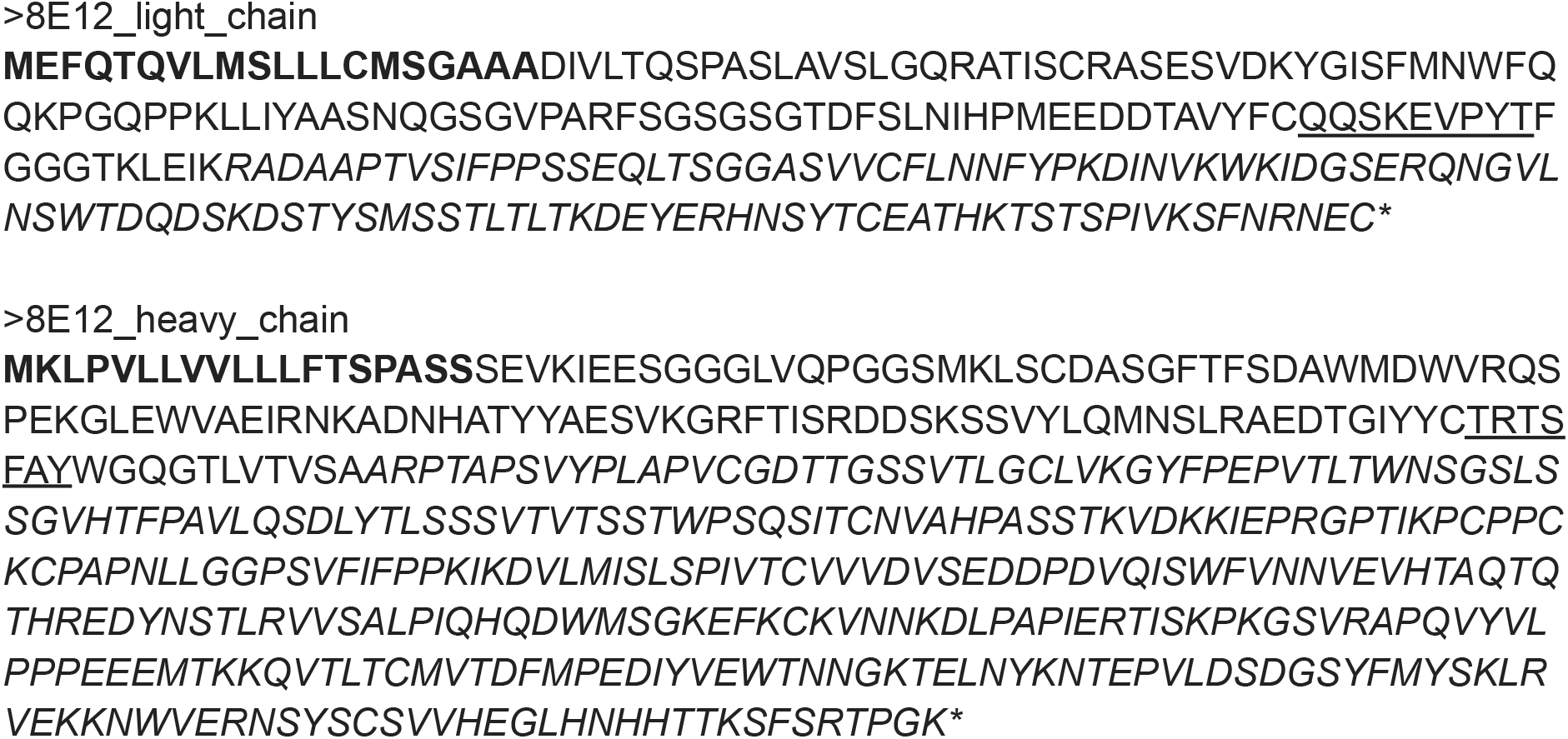
The protein sequences of the anti-IFX 8E12 monoclonal antibody light and heavy chains. Secretion signal peptides are shown in bold, and antibody constant regions in italics. The closest sequence similarity matching to mouse V and J region gene segments were: light chain *Igkv3-2*01* and *Igkj2*01*; heavy chain *Ighv6-6*01* and *Ighj3*01*. Complementarity determining regions in both antibodies are underlined.

**Supplementary Figure 2.**
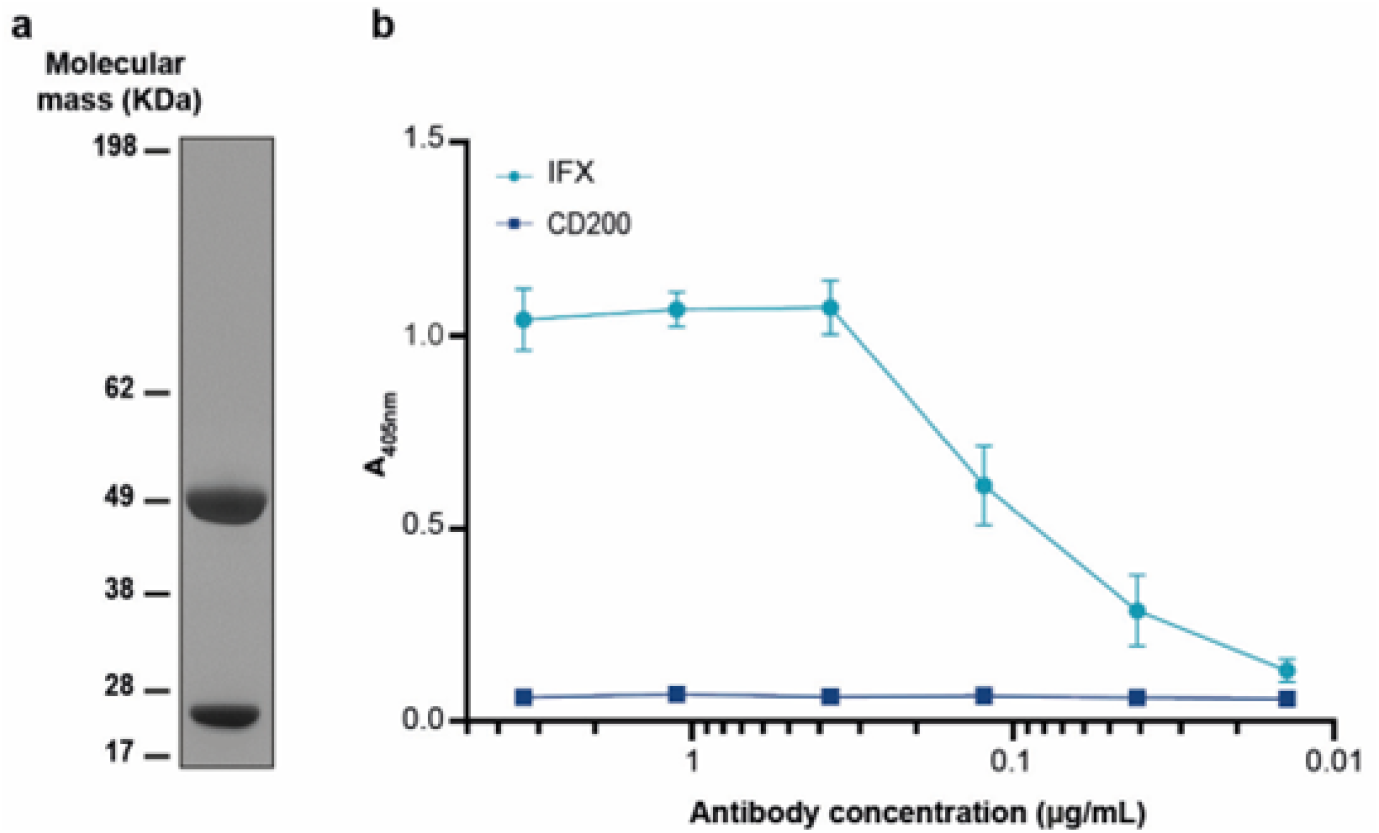
Preparation of a pure and active preparation of the 8E12-IgG2a-formatted monoclonal antibody. HEK293-6E cells were transfected with a single plasmid encoding both light and heavy chains of the 8E12 monoclonal antibody and the expressed antibody purified using protein G resin. (**a**) One microgram of purified antibody was resolved on an SDS-PAGE gel under reducing conditions and shown to be over 95% pure. (**b**) Purified 8E12 antibody is highly active. Purified 8E12 antibody was serially diluted and tested for binding activity to the entire ectodomain of IFX expressed as a soluble enzymatically monobiotinylated recombinant protein captured on a streptavidin-coated microtitre plate; the ectodomain of human CD200 was used a negative control. Data points represent means ± SD; *n*=4.

